# Neonatal infection with *Helicobacter pylori* affects stomach and colon microbiome composition and gene expression in mice

**DOI:** 10.1101/2025.04.30.650706

**Authors:** Katrine B. Graversen, Bella Bjarnov-Nicolau, Sigri Kløve, Krístina Halajová, Sandra B. Andersen

## Abstract

The stomach bacterium *Helicobacter pylori* is estimated to infect half of the world’s population, and the health implications are affected by the age at infection. Neonatal *H. pylori* infection of mice is a relevant model to investigate metabolic and immunological effects. We performed an explorative study at the dynamic first month of life, to compare the composition of the gastrointestinal tract microbiome and stomach gene expression of mice neonatally infected with *H. pylori* with that of uninfected mice. We found that *H. pylori* was present only in the stomach, and that *H. pylori* loads increase with age from one week after infection and onwards, especially after weaning. Stomach and colon microbiome composition was strikingly similar between sites at the same sampling time, but changed significantly over one week, with increased diversity at both sites. Despite that the relative abundance of *H. pylori* in the stomach was low and never exceeded 3%, the composition and alpha diversity of the gastrointestinal microbiome was significantly affected by infection. In a pathway enrichment analysis we found that stomach gene expression related to the extracellular matrix, muscle contraction, and metabolism was affected by infection. Expression of these key processes was, in infected mice, shifted away from that of control mice, towards that of all mice sampled the subsequent week, which we speculate represents accelerated development in infected mice.

## Introduction

The evolutionary history and health implications of infection with the bacterium *Helicobacter pylori* makes it a focus of experimental studies. Specialised on life in the acidic human stomach, *H. pylori* is estimated to infect half of the world’s population, with the association dating back 100-300,000 years (1, 2). It was discovered as the causative agent of stomach ulcers and cancer, where the disease risk is affected by factors such as host-microbe genomics and evolutionary match, and diet (3–5). The immune response to infection affects host development and response to other health challenges (1). In correlative human studies, *A. H. pylori* infection is associated with a lower risk of asthma and allergies (2–4). Mechanistic mouse experiments show that early-life infection has systemic anti-inflammatory effects through direct interaction with dendritic cells, promoting tolerance that can protect against disease (5, 6). The gastrointestinal microbiota also affects immune system development and perturbations to the composition, particularly early in life, can cause lasting health effects. While *H. pylori* infection has been shown to affect the composition of the gastrointestinal microbiome in humans and mice (7–11), the effects of infection early in life in particular are not well characterized.

The environment of the gastrointestinal tract is highly dynamic in the early life of mammals, and provides signals essential for immune system development (12). Sterile at birth, the tract is colonised by microbes from the mother and the environment, and the initial milk diet contributes maternal immune components, nutrients, and microbes that can metabolise them (13–15). Diversity in the intestinal microbiome increases as variety is introduced in the diet, and stabilises in humans at an adult level around the age of three years (16) and in mice at three to four weeks (17, 18). This triggers a strong immune response that facilitates appropriate future inflammatory responses (19). Transmission of *H. pylori* is primarily within families (20, 21), assumed via a gastro-oral or gastro-fecal route, and takes place in the first years of life where stomach pH and immune tolerance are higher (22). *H. pylori* has a range of adaptations to avoid the gastric acid, allowing it to thrive in the stomach niche (23). In contrast, it cannot compete against other microbes in the lower intestine, and in infected adults *H. pylori* DNA can only rarely be detected in feces (24–26). When mice are infected with *H. pylori* in the first week of life, bacterial densities are initially low followed by a high stable load after weaning with low grades of inflammation (5, 27). This is in contrast to adult infection, where inflammation is more extensive and final loads lower (5, 27). As we believe early-life *H. pylori* infection of mice is a relevant model to investigate metabolic and immunological effects resembling human infection, we performed an explorative study, to compare the gastrointestinal tract of mice in the first month of neonatal infection with *H. pylori* with that of uninfected individuals. We show that bacterial loads increase with age as expected and found that *H. pylori* cannot be detected in the lower intestinal tract early in infection. Despite low bacterial loads in the two weeks following *H. pylori* infection, the composition of the gastrointestinal microbiome was affected. Stomach gene expression related to the extracellular matrix, muscle contraction, and metabolism was altered in a way we speculate represent accelerated development.

## Methods

### H. pylori culture

*A. H. pylori* inoculum for mouse infection was prepared as previously described (28). In short, a frozen stock of *H. pylori* strain PMSS1 was grown on TSA Sheep Blood plates (Thermo Fisher Scientific) for 72h in microaerobic conditions. Colonies were swabbed into Brucella broth (BD Biosciences) with 10% fetal bovine serum (FBS, Sigma-Aldrich) and 0.06 mg/ml vancomycin (Sigma-Aldrich), and grown overnight with 100 rpm shaking. Cultures were diluted a few hours prior to use, and motility of bacteria checked in the microscope. The liquid culture was centrifuged at 2000 rpm for 10 min and the pellet resuspended in fresh Brucella broth with 10% FBS. The culture was plated in serial dilutions on TSA Sheep Blood plates to estimate CFU after 72h growth, which was 2.4*10^9^ - 1.5*10^11^ CFU/mL.

### Animal experiment outline

Neonatal C57BL6/J mice were orally gavaged at 6 and 7 days of age with 50 ul of *H. pylori* PMSS1 culture (30 mice, 5 litters in 3 cages), or growth media alone as control (12 mice, 2 litters in the same cage). Groups of 6 mice were sacrificed two days, or one, two, three or four weeks after gavage. We had fewer mice in the control group, and these were divided between the week one and week two time points (**Table S1**). At sacrifice, stomach and gut were dissected. The empty stomach was divided into the upper and lower part, approximately constituting the forestomach and the glandular part of the stomach. Both were cut into halves longitudinally, and one part was immediately cut into small pieces. All four stomach samples and approximately 1 cm pieces of small intestine and colon with content were snap-frozen on dry ice and stored at -70 °C. The animal experiment was performed on license 2020-15-0201- 00528.

### DNA extraction

Total DNA was extracted from one piece of the upper and lower stomach and small intestine tissue of *H. pylori* infected mice, and the lower stomach of control mice from week one and two after gavage, using the DNeasy Blood & Tissue Kit (Qiagen, Germany). The tissue was weighed, cut into small pieces, and put into 2 ml Eppendorf tubes together with 180 ul ATL lysis buffer and one sterile aluminum bead. The tissue was mechanically lysed using the Qiagen TissueLyser for 40 seconds at 15 Hz. Samples were incubated with proteinase K at 56 LJ for 2 hours with rotation and DNA was extracted according to the manufacturer’s protocol. DNA was extracted from colon content from mice week one and two after gavage, with the Zymo Quick-DNA Fecal/Soil Microbe 96 Magbead Kit (Zymo Research, USA) according to the manufacturer’s protocol. A blank extraction control was included for stomach and colon samples.

### *H. pylori* quantification

For samples of infected mice, copy numbers of *H. pylori* from lower and upper stomach sections and small intestine were estimated by qPCR targeting the single-copy *H. pylori* gene glmM (28), in samples from day 2 - 4 weeks after gavage. A few control mice were included as negative controls. We used mastermix from qPCR BioSygreen (PCRBioSystems), and absolute copy numbers were calculated using a standard curve based on amplicons of diluted plasmid containing the glmM gene sequence (28). Absolute copy numbers were standardised to tissue weight. We tested whether there was a change in density over time in weeks with a linear model with location (upper or lower stomach) as a fixed effect.

### Microbiome analyses

The V3-4 region of the 16S rRNA gene was sequenced from lower stomach tissue and colon content, from mice sacrificed one and two weeks after gavage. Library preparation and paired-end Illumina NovaSeq 6000 sequencing was performed at Novogene UK. Microbiome analyses were performed in RStudio (ver. 2023.12.1.402, (29)). The in-silico decontamination tool SCRuB (30) was used to remove reads classified as contamination based on the extraction controls. Reads were processed with the DADA2 pipeline (31). The reads were trimmed with truncLen = c(220,220) and merged with an overlap of 10 bp. Chimeras were removed and taxonomy assigned using the Silva reference database v138.1 (32). In phyloseq (33) we removed chloroplast, mitochondria, and eukaryote sequences and pruned ASVs that occurred less than 10 times. ASVs were agglomerated using the tip_glom function with h = 0.03. phyloseq was also used for calculating the alpha diversity measured as the Shannon diversity index. The differences between groups at the different timepoints were determined by two-way analysis of variance (ANOVA) and Tukey’s post hoc test. Beta diversity was measured by the weighted UniFrac distance matrix and statistical differences were calculated with the permutational multivariate analysis of variance (PERMANOVA) test in vegan (34). The MicroViz package was used to make the compositional barplot (35).

Differentially present bacterial genera were identified with LeFSe using microbiomeMarker (36).

### RNA extraction and sequencing

RNA was extracted from lower stomach tissue samples from week one and two after gavage with Qiagen MiRNeasy Tissue/Cells Advanced Mini Kit according to the manufacturer’s protocol. With a sterile aluminum bead, tissue was disrupted and homogenized in the Qiagen TissueLyser for four minutes at 25 Hz and an additional DNase treatment was included.

Samples were sequenced in two rounds on Illumina NovaSeq 6000, at the Department of Genomic Medicine, Rigshospitalet, Copenhagen, Denmark with TruSeq Stranded Total RNA library preparation, and at Novogene UK targeting all mRNA.

### Gene expression analysis

The RNA-sequencing data was trimmed to remove poly-G tails (--trim_poly_g) and filtered to remove duplicate reads with maximum accuracy (dup_calc_accuracy = 6) by fastP (37). It was processed using the nf-core/RNA-seq pipeline v3.14.0 (38) with the default settings, aligning the reads to the provided mouse reference genome GRCm39 release no. 109 using STAR (39) and quantifying mapped reads using Salmon (40).

Principal coordinate analysis (PCA) and differential gene expression analysis was performed on a gene length scaled count matrix with DESeq2 (41) in RStudio (ver. 2023.12.1.402, (29)), calculated using Wald statistics with Bonferoni corrections for multiple testing. Gene counts were first rounded to integers and the count matrix was filtered to genes with counts > 0, that were present in > 10 samples. Two designs were applied in the analysis. To extract DE genes between *H. pylori* infected and controls across both timepoints, included treatment, time, and sequencing run (design = ∼ Treatment + Time + Run) with contrast = c(“Treatment”, “Hp”, “Ctrl”). To extract DE genes with p_adj_ < 0.05 at each of the individual time points, a combined four-level factor consisting of treatment and time was included in the design (design = ∼ Treatment_Time + Run) with contrast = c("Treatment_Time", "Hp_W1", "Ctrl_W1") and contrast = c("Treatment_Time", "Hp_W2", "Ctrl_W2"). Statistical differences between treatments, sampling time, and sequencing run on overall gene expression were calculated by PERMANOVA test in vegan::adonis2 (34) based on Bray– Curtis dissimilarity matrix between samples.

To identify enriched pathways, we applied STRING (version 12.0; https://string-db.org/; (42)) on differentially expressed genes with a more permissive cutoff of p_adj_ < 0.1 from the DESeq2 results, for week one, week two and across both weeks. We applied STRING to total gene lists. As background, we used all the genes with detected expression. Within STRING we chose to report Reactome pathways (43) with a signal > 1. The signal represents a weighted harmonic mean between the observed/expected ratio and -log(FalseDiscoveryRate).

## Results

### *H. pylori* colonisation

In stomachs, there was an initial drop in *H. pylori* loads from day 2 to one week post gavage, likely representing the wash-out of the initial inoculum (**Fig. 1**) . From week one after gavage and onwards, there was a significant increase in *H. pylori* counts adjusted for tissue weight (**Fig. 1**; Linear regression, F = 10.26, df = 3, 44; R^2^_adj_ = 0.37; p_adj_ < 0.001). There was a non-significant trend for higher loads in the lower stomach compared to upper stomach (Tissue location: t = -1.91; p = 0.062). No *H. pylori* was detected in any small intestine samples (data not shown).

**Fig. 1:**
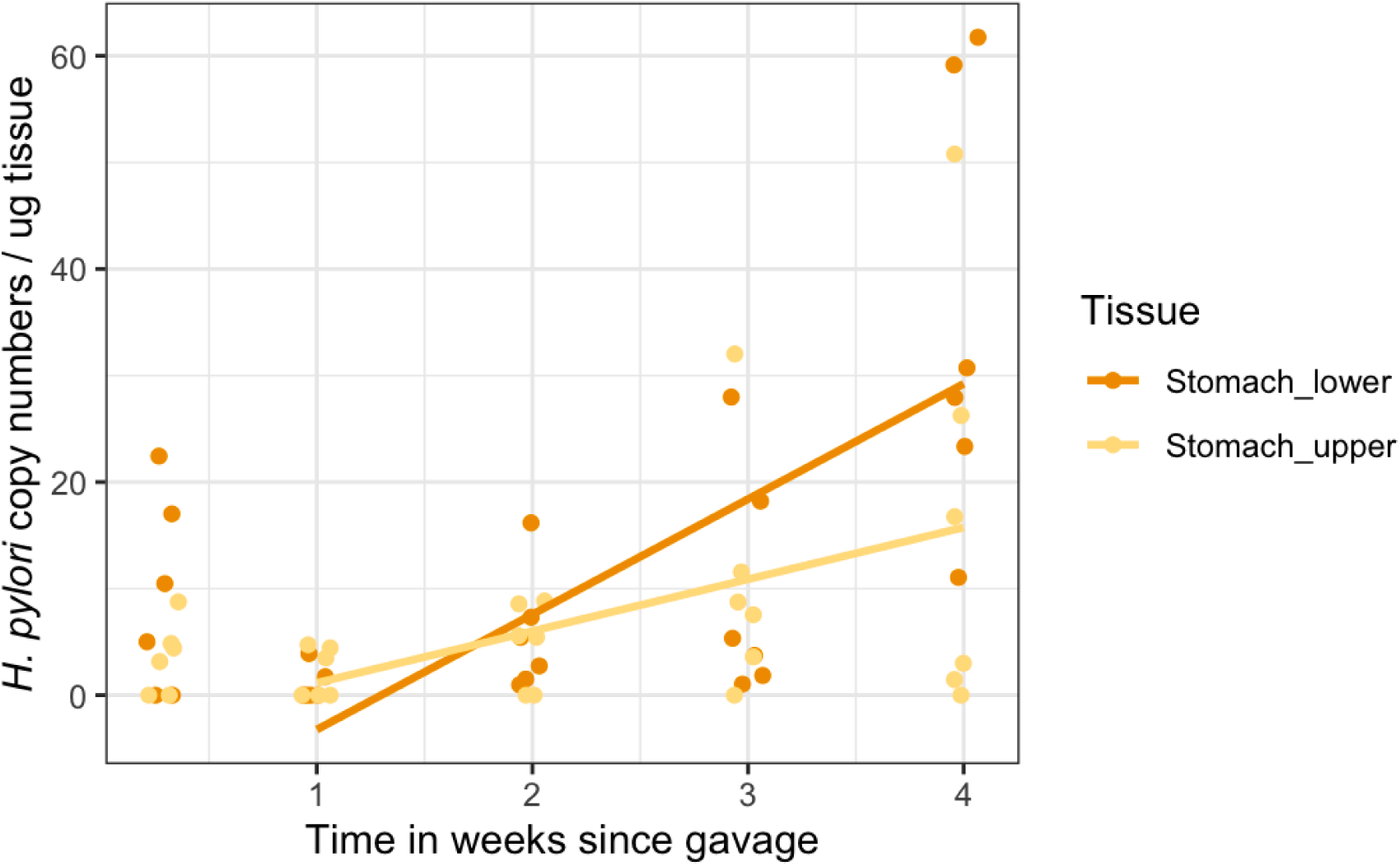
The H. pylori copy numbers per ug tissue detected by qPCR increases significantly over time from week one to four weeks after gavage. There was a trend towards a greater increase in bacterial load in the lower part of the stomach. From two days to one week after gavage there was a drop in bacterial load, which likely represents wash-out of the inoculum.

We visualised the microbiome composition in stomach and colons sampled one and two weeks after gavage, by depicting the relative abundances of the top 15 most abundant taxa at genus level. Strikingly, the stomach and colon samples from the same week are more similar to each other, than to those from the same site collected the other week (**Fig. 2**). Samples from week one are dominated by *Ligilactobacillus* lactic acid bacteria, while a Muribaculaceae genus expands in week two. In week one, the LEfSe analyses showed *H. pylori* to be enriched in the stomach of infected mice, and *Enterobacter* in the colon, which was specific to litter 15. In the control mice, Lachnospiraceae Family was enriched in the stomach and *Bacteroides* in the colon in week one (Data not shown). In week two, the *H. pylori* infected mice from litter 15 with *Enterobacter* experienced an additional bloom of *Lachnoclostridium*, which came out as significantly enriched genera in the stomach in the LEfSe analysis (**Fig. 3**). *Akkermansia* was significantly more abundant in both stomach and colon samples in week two in *H. pylori* infected mice (**Fig. 4A**). *Helicobacter* was present only in stomach samples of infected individuals, and even here only reached a median prevalence of 0.11% in week one and 0.64% in week two (**Fig. 4B**). Based on the compositional plots we analysed the alpha and beta diversity for each week separately.

**Fig. 2:**
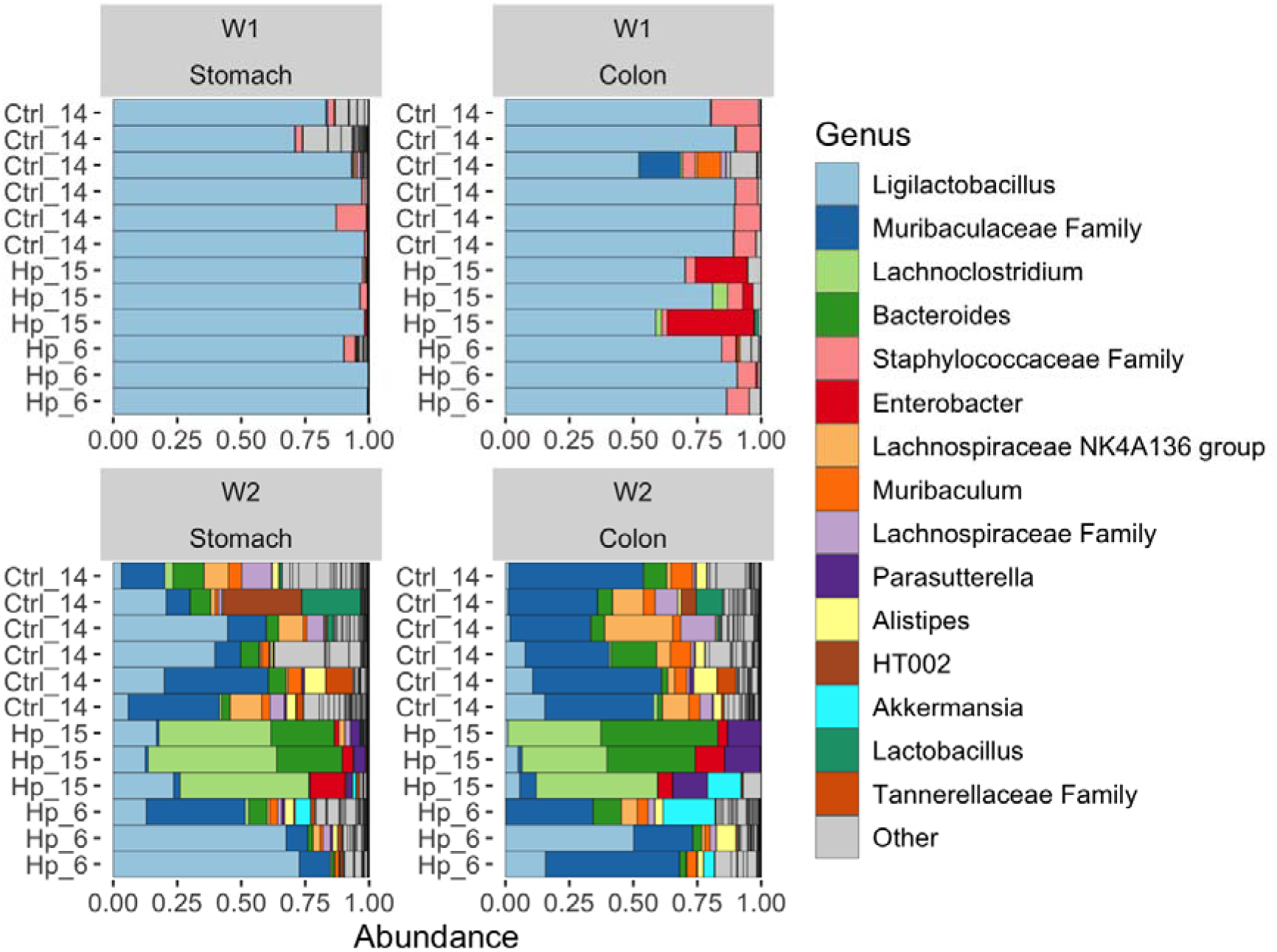
Distribution of the 15 most abundant genera in the stomach and colon samples, sampled one week (W1) and two weeks (W2) after gavage. Samples are denoted by treatment, which is either gavage with a control (Ctrl) solution or H. pylori (Hp) culture, and litter is indicated by number.

**Fig. 3:**
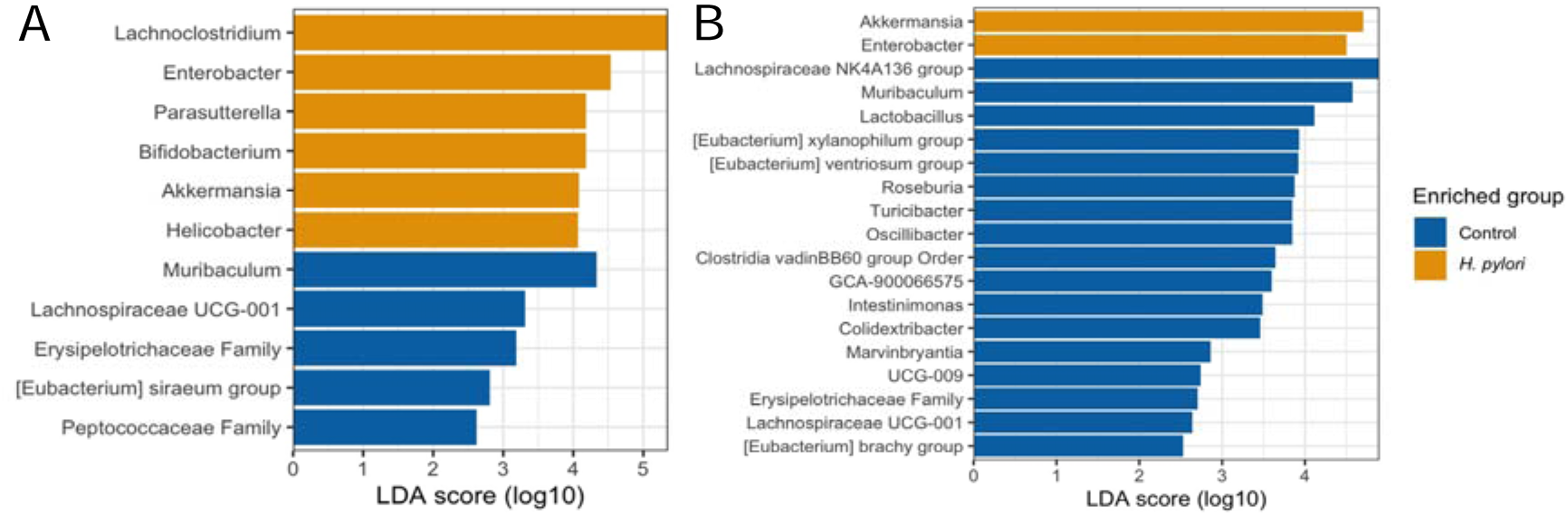
Genera found to be differentially present in control (blue) or H. pylori (yellow) samples A) stomach and B) colon samples from two weeks after gavage by LEfSe. On the x-axis is the effect size score given by Linear Discriminant Analysis.

**Fig. 4:**
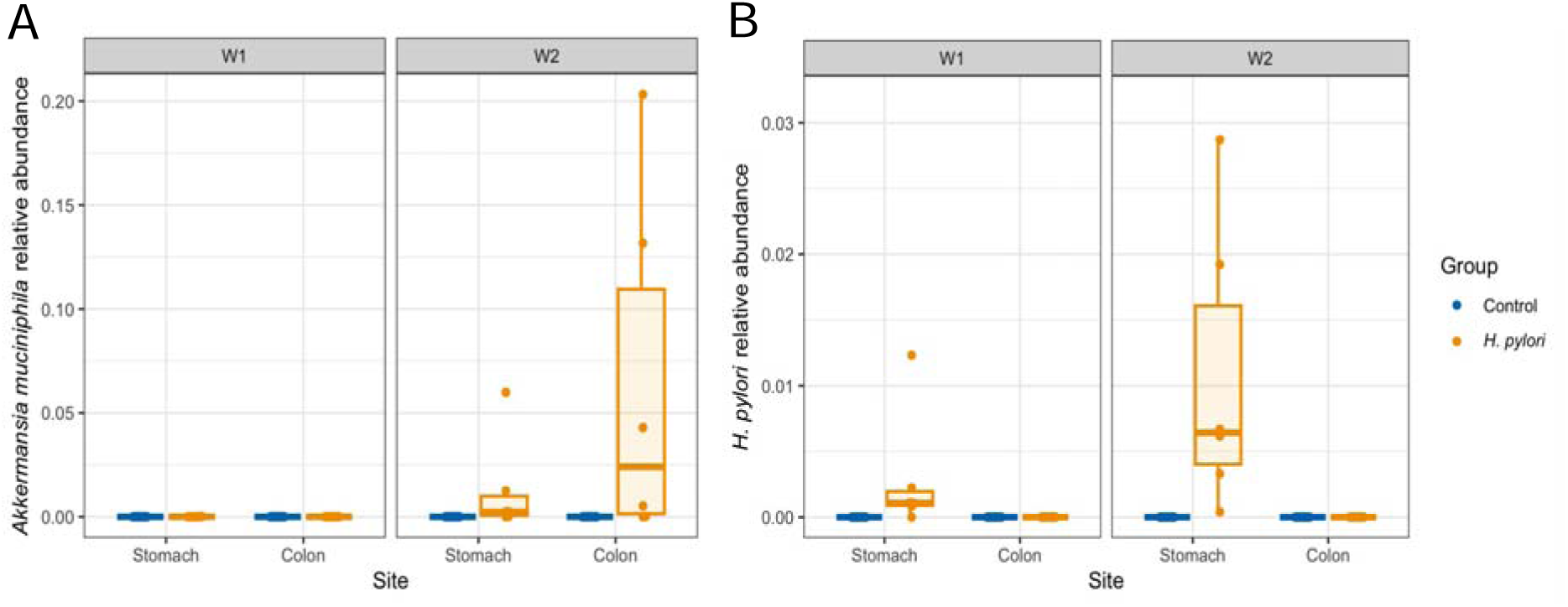
Relative abundance of A) Akkermansia muciniphila and B) Helicobacter pylori in stomach and colon samples from week one (W1) and week two (W2) after gavage.

### *H. pylori* infection lowers alpha diversity

The alpha diversity, measured as the Shannon diversity index, increases from week one to week two (Fig. 5, Table 1). In week one there are no significant differences between treatments but a trend for lower diversity in the stomachs compared to colons for the Shannon index (Table 1). In week two, diversity is lower in samples from *H. pylori* infected mice and significantly lower in the colon of infected mice compared to control colons (p_adj_ = 0.022) with the same trend for the stomach p_adj_ = 0.061).

**Fig. 5.**
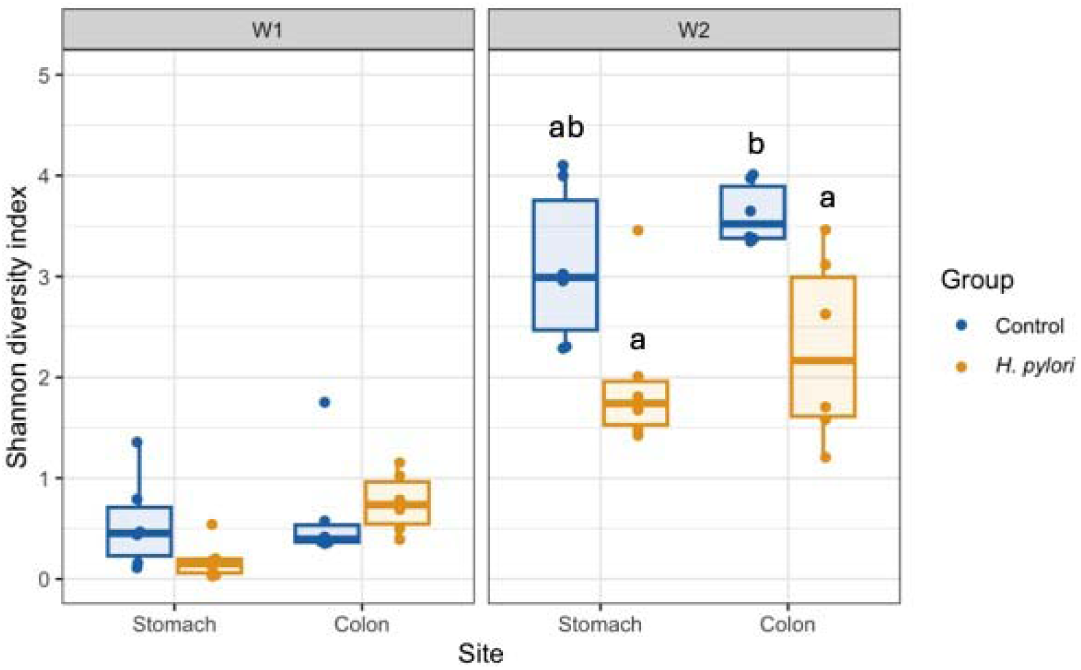
Alpha diversity measured by Shannon diversity index of gastrointestinal samples from early-life H. pylori infected mice compared to controls. Samples were taken from stomachs and colons of mice that were sacrificed one (W1) and two weeks (W2) after gavage. Letters indicate groups that have significantly different means at the 0.05 level following 2-way ANOVA and Tukey HSD. The boxes depict median values and interquartile range, and whiskers the minimum and maximum values, with individual data points as dots.

**Table 1.** Two-Way ANOVA results from alpha diversity (Shannon diversity index) of colon and stomach samples of early-life H. pylori infection in mice. The analyses were performed for week 1 and week 2 alpha diversity independently.

|  | Week 1 |  | Week 2 |  |
| --- | --- | --- | --- | --- |
|  | p-value | F-value | p-value | F-value |
| Group | 0.452 | 0.587 | 0.000484*** | 17.302 |
| Site | 0.059 | 4.013 | 0.184 | 1.892 |
| Group:Site | 0.153 | 2.207 | 0.735 | 0.118 |

### Beta diversity plot shows differences in phylogenetic profiles

To further analyze the differences in the microbial composition of our samples, we looked at sample-to-sample similarities in a beta diversity plot of weighted unifrac distances that relies on phylogenetic distances (**Fig. 6**). In week one there was a significant difference between sites. In week two, there were significant differences between sites and treatments and a trend towards an interaction (**Table 2**).

**Fig. 6:**
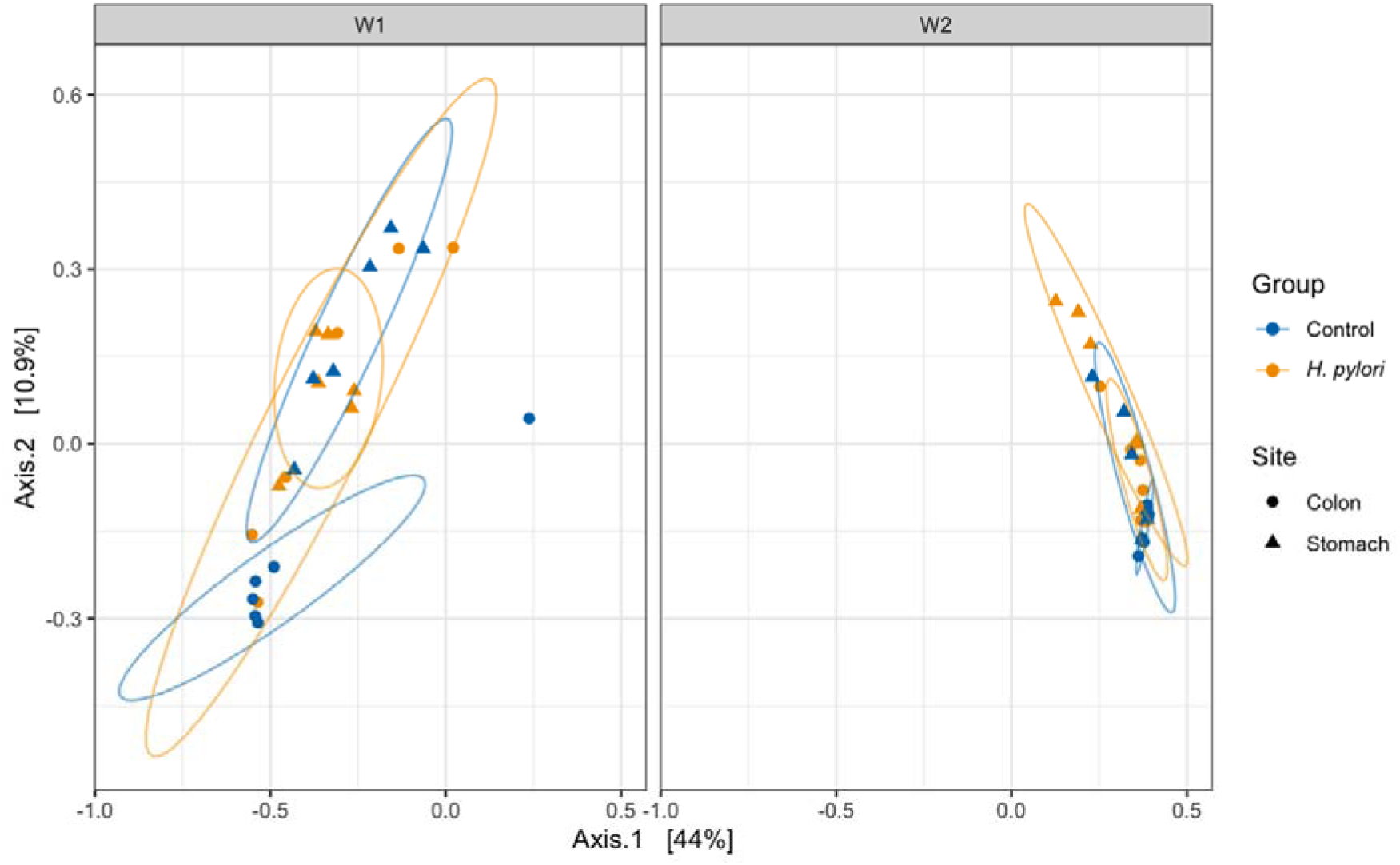
Weighted UniFrac distances depicting beta diversity of colon (circles) and stomach (triangles) samples from early-life H. pylori infection in mice, collected one week (W1) or two weeks (W2) after gavage with H. pylori (yellow) or a control solution (blue).

**Table 2.** PERMANOVA results from beta diversity (Weighted UniFrac distance) measured from colon and stomach samples from early-life H. pylori infection in mice.

|  | Week 1 |  |  | Week 2 |  |  |
| --- | --- | --- | --- | --- | --- | --- |
|  | F-value | p-value | R<sup>2</sup> | F-value | p-value | R<sup>2</sup> |
| Group | 5.3173 | 0.001*** | 0.193 | 4.4913 | 0.001 *** | 0.1583 |
| Site | 0.9048 | 0.544 | 0.0328 | 2.2958 | 0.013 * | 0.08092 |
| Group:Site | 1.3643 | 0.121 | 0.0495 | 1.5841 | 0.078 | 0.05583 |

### Gene expression analysis

A total of 22379 genes were found to be expressed in our stomach tissue samples. A principal component analysis (PCA) of expression profiles from the DESeq2 results show a clear separation between sequencing run (labels) along PC1, with 51% of the variance explained along this axis **(Fig 7**). There was a significant difference between sampling times and sequencing run (**Table 3**). Despite no clear separation by treatment on the PCA plot, the PERMANOVA analysis revealed a borderline significant effect of treatment on overall gene expression when accounting for effects of sampling time and sequencing run

**Fig. 7:**
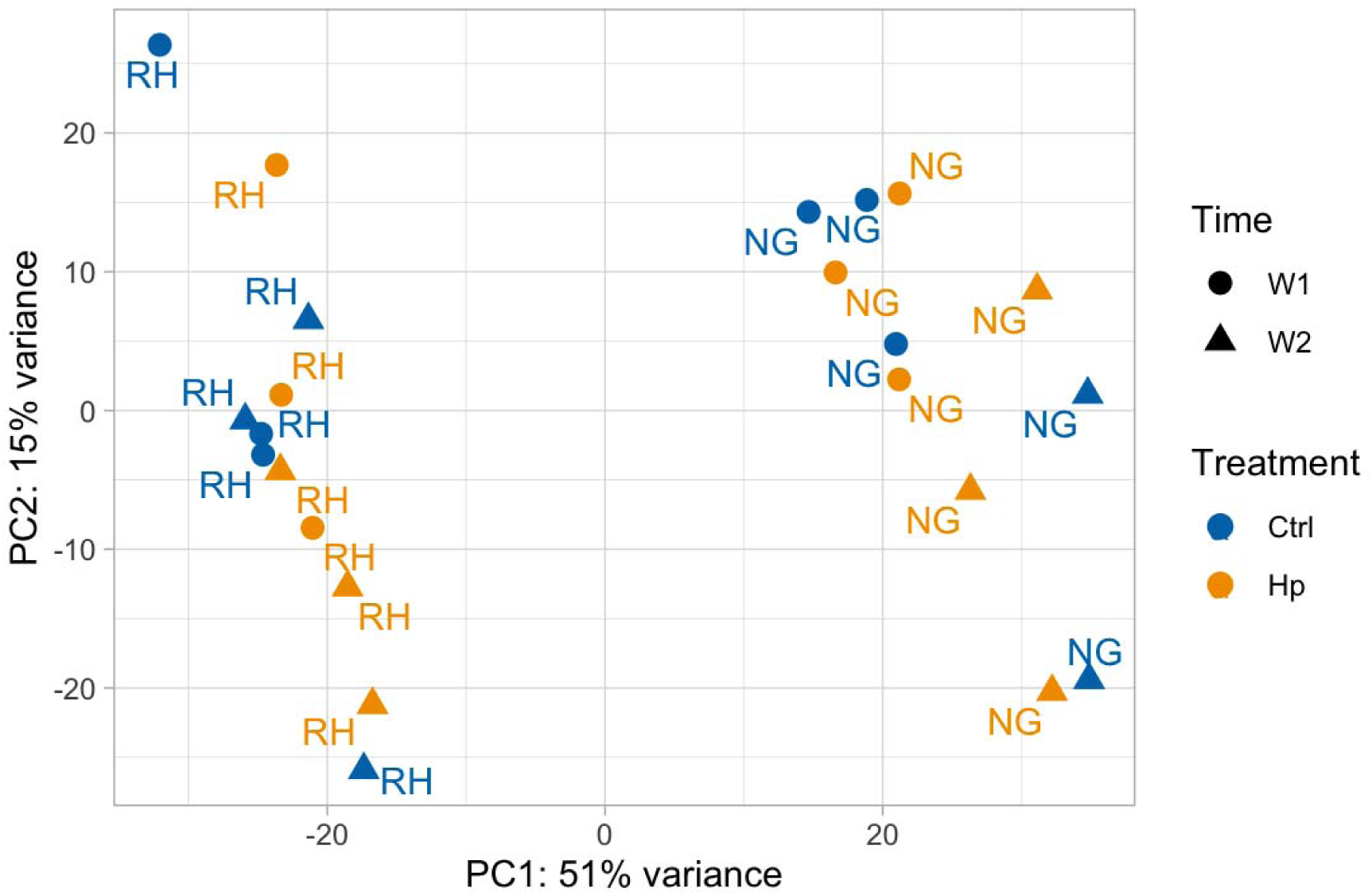
Principal Component Analysis (PCA) plot of stomach RNA-sequencing data from early-life control or H. pylori infected mice illustrating differences in sequencing run (RH = Rigshospitalet, NG = Novogene), time (circles: week one samples, triangles: week two samples), and treatment (blue: control, yellow: H. pylori).

**Table 3.** PERMANOVA results of stomach RNA-sequencing data from early-life control or H. pylori infected mice. No interactions between factors were found to be significant and were thus removed from the model.

|  | F-value | p-value | R<sup>2</sup> |
| --- | --- | --- | --- |
| Treatment | 2.7111 | 0.056 . | 0.02832 |
| Time | 23.1117 | 0.001 *** | 0.24140 |
| Run | 50.9181 | 0.001 *** | 0.53183 |

To identify differentially expressed genes between treatment groups, DESeq2 analysis was performed with two designs, to analyze the effect across both timepoints and at the individual time points, controlling for sequencing run. In total, only 13 genes were found to be significantly differentially expressed (p_adj_ < 0.05); four genes by treatment across both timepoints (*Gpt, Magix, Mns1* and *Shisa3*); three genes when performing the analysis on only week one data (*Ddr, Itpr1,* and *Tnni3*) and six genes when performing the analysis on week two data (*Dusp1, Exo5, Reg3g, Siah2, Csn2* and *AI197445*). The overall pattern of expression between groups was the same in the two runs, however, values were generally lower in the Novogene data set (**Fig. S1**). This was accounted for by controlling for sequencing run in the DEseq2 analyses.

### STRING network revealing enriched pathways

To identify enriched pathways, we increased the cutoff to a more permissive p_adj_< 0.1 and extracted the DE genes for week 1 (68 up, 116 down), week 2 (5 up, 12 down) and across timepoints (13 up and 11 down). We found enriched Reactome pathways at a signal cut-off of 1 for genes that were differentially expressed in week one after gavage (**Fig. 8** & **9**). For these genes, the expression in *H. pylori* infected mice in week one resembled that of all mice sampled in week two more than that of the control mice in week one (**Fig. 10**). In the upregulated set, 12 genes were classified in the “Citric acid cycle and respiratory electron transport” pathway, and subsets of these genes were further classified as various pathways related to Complex 1 biogenesis and respiratory electron transport (**Fig. 8**). These and an additional 19 genes were also classified in a broad “Metabolism” pathway. For the majority of these genes, expression was lowest in week one, with tissue from *H. pylori* infected mice exhibiting higher expression than that from control mice, and with expression higher and similar between groups in week 2 (**Fig. 10**).

**Fig. 8:**
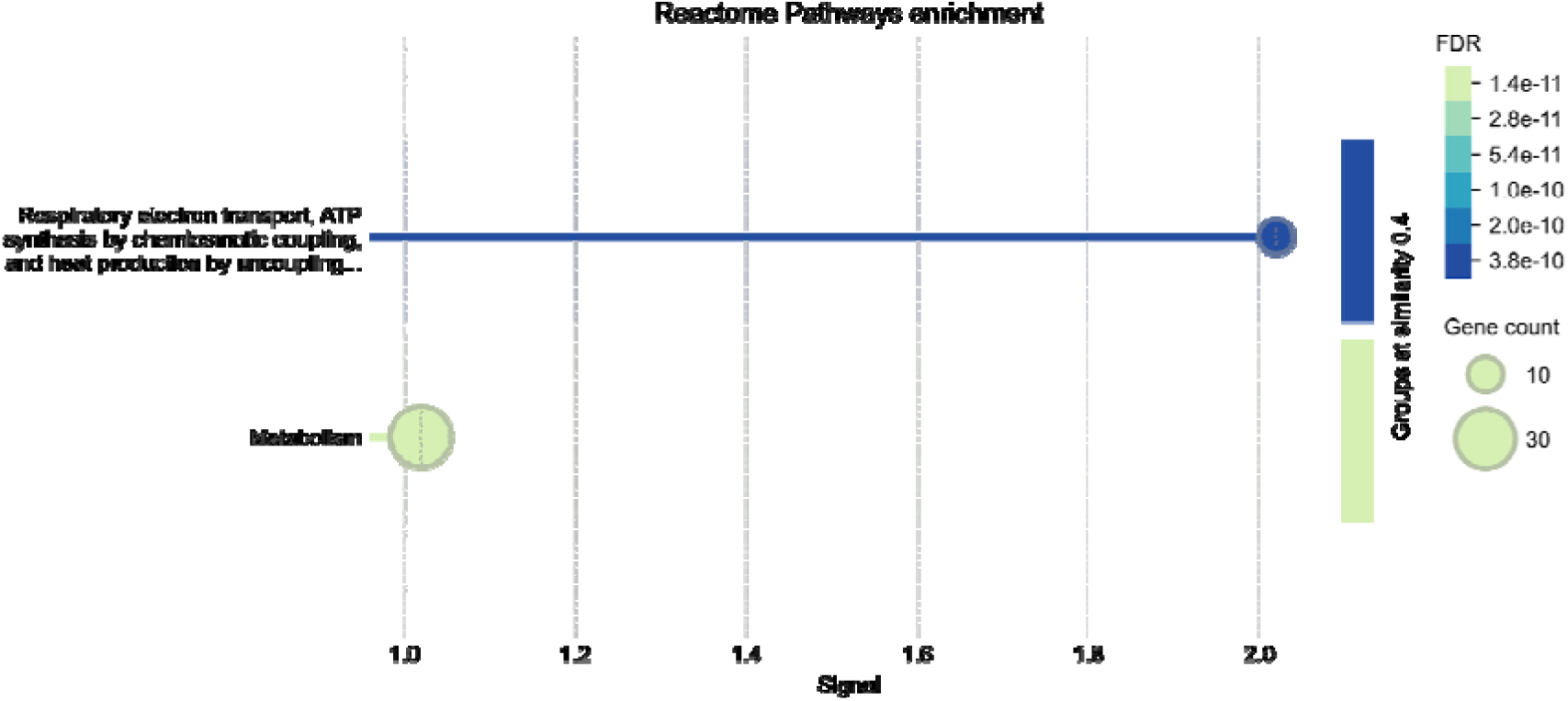
Enriched pathways of upregulated genes (padj < 0.1) in week one samples from H. pylori infected mice. Illustration from STRING for signal > 1 and merging terms with a similarity ≥ 0.5.

In the set of genes downregulated by *H. pylori* infection, 18 genes were classified in the “extracellular matrix organization” pathway (including the *Ddr* gene with p_adj_ < 0.05) and subsets of these genes were further classified as various pathways related to collagen (**Fig. 9**). An additional 10 genes (including *Itpr1* with p_adj_ > 0.05) were classified in the Muscle Contraction pathway. While genes related to muscle contraction were generally downregulated, two upregulated genes also belonged to this pathway (including *Tnni3* with p_adj_ > 0.05). For the majority of the genes, expression in tissue from control mice was higher in week 1 compared to week 2, and infection with *H. pylori* lowered expression in week 1 to resemble the expression of all mice in week 2 (**Fig. 10**).

**Fig. 9:**
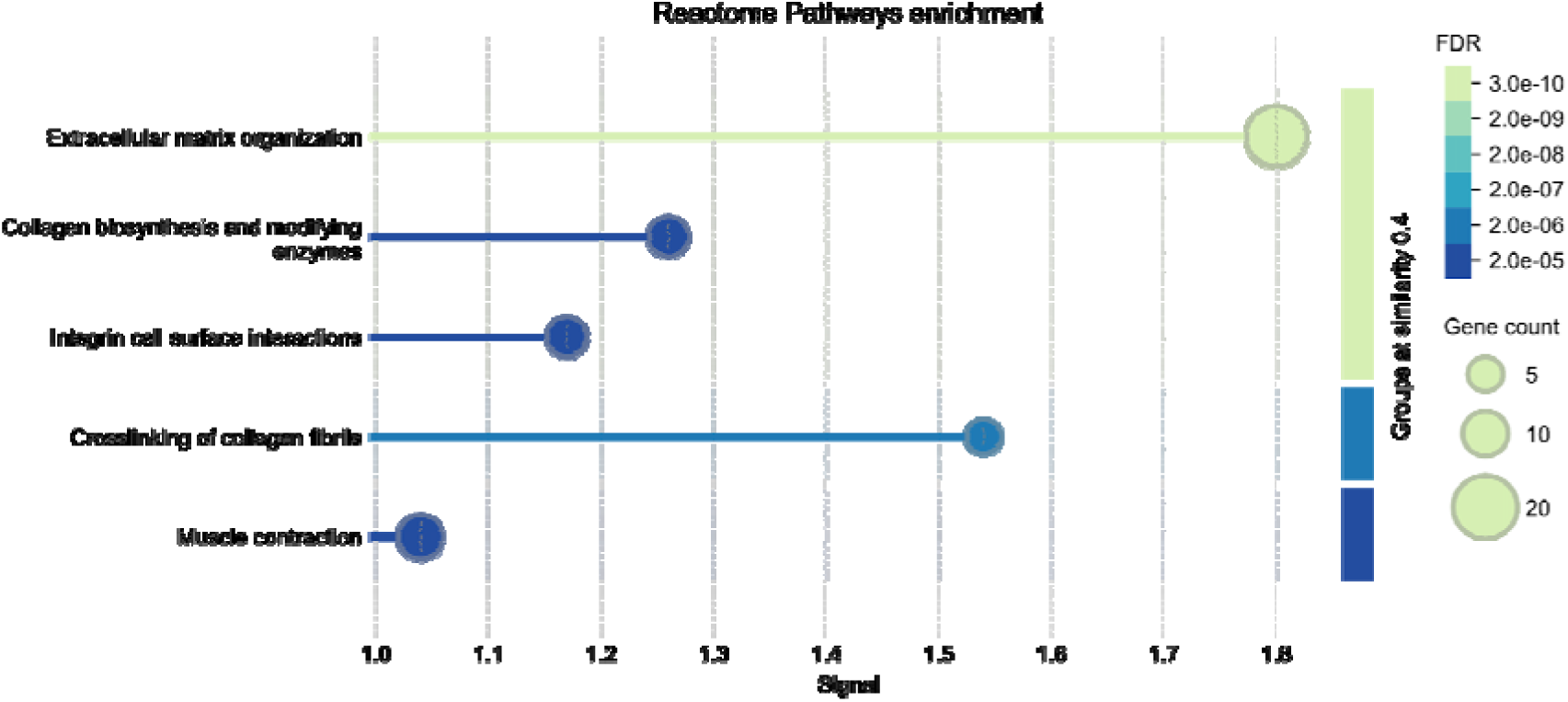
Enriched pathways of downregulated genes (padj < 0.1) in week one samples from H. pylori infected mice. Illustration from STRING for signal > 1 and merging terms with a similarity ≥ 0.5.

**Fig. 10:**
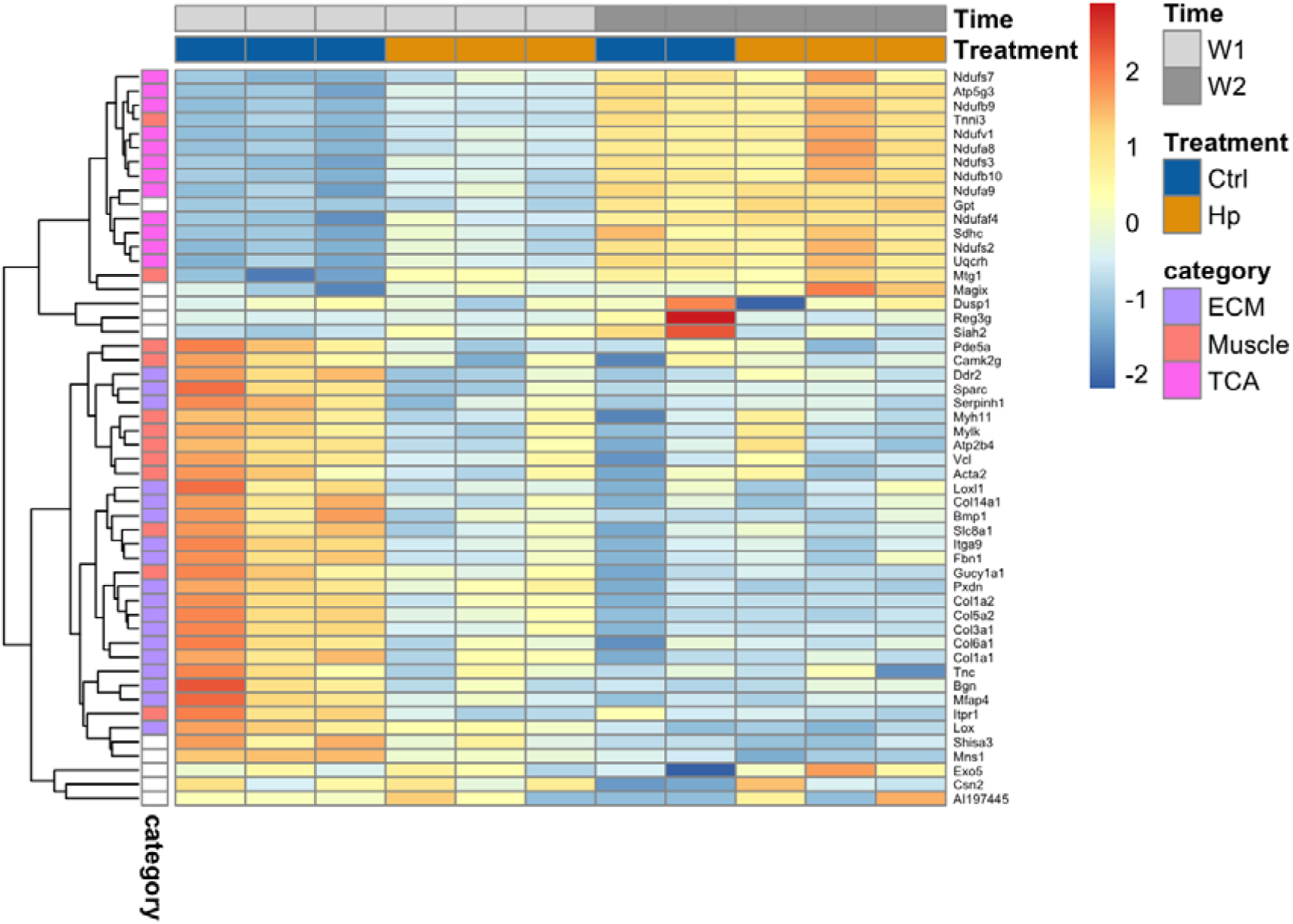
Heatmap of scaled expression of genes that were differentially expressed with a padj < 0.05, or 0.1 and classified as belonging to an enriched Reactome pathway in the STRING analyses (ECM = extracellular matrix, Muscle = muscle contraction, and TCA = the citric acid cycle). Only data from Novogene is shown, due to the differences in expression levels between sequencing runs. See Fig. S2 for transcriptomic data from RH showing the same patterns.

## Discussion

### *H. pylori* colonises the entire stomach in the first month of life

We characterise the impact of experimental *H. pylori* infection on the mouse gastrointestinal environment, at the dynamic first month of life. Stomach morphology and gene expression change radically at this time, as the mice gradually start eating solids around two weeks of age (44). We found that the initially low *H. pylori* load increased, with a trend towards a preference for the lower glandular part of the stomach, as previously found (5, 45, 46) (**Fig. 1**). The upper part that is the forestomach does not contain the glands that house *H. pylori* (45, 46), which fully develop around 3 weeks after gestation (47). In the small tissue sections, we did not distinguish between the antrum and corpus part of the lower glandular stomach, where *H. pylori* early in infection has been found to reach higher loads in the antrum (46). The relative abundance of *H. pylori* in the stomach one and two weeks after gavage was low and never exceeded 3%. This is in contrast to mean levels at 26% in adult mice following the same gavage treatment early in life (28). We did not detect the bacteria in the small intestine by qPCR or in the colon by 16S rDNA sequencing (**Fig. 4**). The gastric pH in young mice is higher than that of adult mice (around pH 4-5 at one week compared to < 3 (48) or around 3 (49) in adults. The higher gastric pH reduces the competitive edge in the stomach that acid tolerance provides *H. pylori* with. While the neonatal immune system on one hand is tolerant of microbial associates (12), maternal antibodies and milk oligosaccharides may bind and limit bacterial growth (15).

### *H. pylori* infection shape the stomach and colon microbiomes despite a low relative abundance

The stomach and colon microbiomes were strikingly similar between samples from the same week and alpha diversity was low. One week after gavage, the mice were two weeks old and primarily nursing, which is reflected in the microbiome dominated by lactic acid bacteria. There has been limited research done on neonatal stomach microbiomes, one study found *Lactobacillus* dominance but did not compare with the gut microbiome (50).

*Ligilactobacillus* has previously been found to dominate the intestinal microbiome in neonatal mice, driven by the uniform milk diet, and *Staphylococcaceae* genera have likewise been sampled from neonatal mice likely originating from maternal skin, oral or vaginal sites (17). The similarity between stomach and colon microbiomes week one might be related to the higher gastric pH in neonatal mice making the conditions along the gastrointestinal tract more uniform. The alpha diversity rose drastically between week one and week two (**Fig. 2**). The gastrointestinal microbiome is developing fast in this early stage of life, and two weeks after gavage, the mice are ingesting more solid food. At this time mice also become coprophagic, which means that they ingest their own and nest mates’ stool. Thus, the detected bacteria are not necessarily colonizing the stomach but might just be ingested, and this could in part explain the observed rise in diversity of the stomach microbiome, and its similarity to that of the colon.

Two weeks after infection, alpha diversity was significantly reduced in colon samples from *H. pylori* infected mice and a similar trend was observed for stomach samples (**Fig. 5**), despite *H. pylori* only reaching a median prevalence of 0.64% in week two (**Fig. 4B**). The lower alpha diversity of *H. pylori* infected mice was thus not driven by a high prevalence of *H. pylori* in these samples, as has been found in humans (7, 9). This points to *H. pylori* being able to alter the diversity in both the colon and the stomach indirectly. In a study where mice were infected with *H. pylori* at 4-6 weeks of age, beta diversity in the ileum and cecum was significantly altered between the infected and control groups (8). The indirect effects are hypothesized to occur through modulation of the immune response, affecting the microbiome composition (8). Pups from the *H. pylori* infected litter 15 had a large proportion of bacteria belonging to *Lachnoclostridium* and *Bacteroides,* causing low alpha diversity. Pups from the same litter had a higher prevalence of *Enterobacter* in week one colon samples that may represent a disturbance of the microbiota.

*A. muciniphila* bacteria reached significantly higher relative abundances in both the stomach and colons of *H. pylori* infected mice (**Fig 4A**). In our recent study of early-life *H. pylori* infection in mice also found a significant bloom of *Akkermansia muciniphila* in *H. pylori* infected mice on a high fat diet, when present in the maternal or littermate microbiome (28). Here we do not know the *A. muciniphila* status of the dams. *A. muciniphila* is mucus degrading and has gained attention as a probiotic, as studies find that it can contribute to weight loss (51–53), and improve the efficacy of immunotherapy in some cancer treatments (54). However, it may also act as an opportunist when the gut microbiota has been perturbed e.g. by antibiotics (55, 56), and we found its abundance to be highest in the *H. pylori* infected mice that had a worse metabolic profile compared to controls (28).

### Gene expression of key stomach development processes affected early in infection

Genes in pathways related to extracellular matrix formation, smooth muscle contraction and metabolism were differentially expressed in *H. pylori* infected mice one week after gavage. These processes have previously been found to be affected by *H. pylori* infection in cell and animal models, and in human biopsies (57, 58). The extracellular matrix provides structure to the stomach tissue, and serves as a medium for intercellular communication and a scaffold for the stomach muscles that facilitate gastric motility. Tissue invasion and disruption by *H. pylori* can expose the underlying extracellular matrix, and the bacteria adhere to components of this structure such as collagen (59). In rats, *H. pylori* infection causes increased collagen deposition that invades the submucosa muscle fibers, and a decrease in muscle contraction force and rate (60). The associated stomach immobility and delayed stomach emptying, has been implicated in patients experiencing dyspepsia (61), while gastric emptying was accelerated in one study of *H. pylori* infection in children (62).

Expression of these key processes was, in infected mice, shifted away from that of control mice, towards that of all mice sampled the subsequent week (**Fig. 10**). The stronger signal of infection on expression in week 1 may be caused by the high bacterial load in the first week from the gavage inoculum (**Fig. 1**), or represent the acute response to infection. The two time points we sampled have been characterised as distinct phases of stomach development, marked specifically by a switch from high expression of focal adhesion structures and epithelial receptors targeting the ECM, towards increased metabolism and maturation (44). Whether the altered expression we observed in infected mice compared to controls, resembling that of one week older mice, reflect faster tissue development, or impairment of growth is unclear. Additional time points and histological data is needed to explore this. The carcinogenic effects of *H. pylori* involves cell proliferation by inducing stemness properties (63). Tumorigenesis of diffuse gastric cancer was found to share cell proliferation markers of early stomach development (44), and the expression differences we observe may reflect brief early reprogramming, with life long effects.

### Strengths and limitations of experimental design

By sampling at multiple timepoints, we were able to capture dynamic changes in the microbiome and gene expression over time, enhancing our understanding of the impact of early-life *H. pylori* infection on host development. The stomach microbiota of mice has been studied less than fecal microbiomes and especially that of neonates (50). Transcriptomics were performed in two batches with different approaches, which introduced extensive technical variation. This likely made it more difficult to detect more subtle differences in expression between groups. Additional time points for the microbiome and gene expression analyses, in combination with histology, would provide more insight into how infection shapes development. We further observed litter effects in the microbiome analyses within the *H. pylori* group and ideally samples from more litters in both groups had been analysed.

### Data availability

Microbiome and RNA sequencing data is available as NCBI BioProject PRJNA1219814.

## References

1. Arnold IC, Hitzler I, Müller A. 2012. The immunomodulatory properties of *Helicobacter pylori* confer protection against allergic and chronic inflammatory disorders. Front Cell Infect Microbiol 2:10.

2. Chen Y, Blaser MJ. 2007. Inverse associations of *Helicobacter pylori* with asthma and allergy. Arch Intern Med 167:821–827.

3. Chen Y, Blaser MJ. 2008. *Helicobacter pylori* colonization Is inversely associated with childhood asthma. J Infect Dis 198:553–560.

4. Reibman J, Marmor M, Filner J, Fernandez-Beros ME, Rogers L, Perez-Perez GI, Blaser MJ. 2008. Asthma is inversely associated with *Helicobacter pylori* status in an urban population. PLoS One 3:3–8.

5. Arnold IC, Lee JY, Amieva MR, Roers A, Flavell RA, Sparwasser T, Müller A. 2011. Tolerance rather than immunity protects from *Helicobacter pylori*–induced gastric preneoplasia. Gastroenterology 140:199–209.e8.

6. Arnold IC, Dehzad N, Reuter S, Martin H, Becher B, Taube C, Müller A. 2011. *Helicobacter pylori* infection prevents allergic asthma in mouse models through the induction of regulatory T cells. J Clin Invest 121:3088–3093.

7. Bik EM, Eckburg PB, Gill SR, Nelson KE, Purdom EA, Francois F, Perez-Perez G, Blaser MJ, Relman DA. 2006. Molecular analysis of the bacterial microbiota in the human stomach. Proc Natl Acad Sci U S A 103:732–737.

8. Kienesberger S, Cox LM, Livanos A, Zhang X-S, Chung J, Perez-Perez GI, Gorkiewicz G, Zechner EL, Blaser MJ. 2016. Gastric *Helicobacter pylori* infection affects local and distant microbial populations and host responses. Cell Rep 10.1016/j.celrep.2016.01.017.

9. Sugihartono T, Fauzia KA, Miftahussurur M, Waskito LA, Rejeki PS, I’tishom R, Alfaray RI, Doohan D, Amalia R, Savitri CMA, Rezkitha YAA, Akada J, Matsumoto T, Yamaoka Y. 2022. Analysis of gastric microbiota and *Helicobacter pylori* infection in gastroesophageal reflux disease. Gut Pathog 14:38.

10. Mannion A, Sheh A, Shen Z, Dzink-Fox J, Piazuelo MB, Wilson KT, Peek R, Fox JG. Jan-Dec 2023. Shotgun metagenomics of gastric biopsies reveals compositional and functional microbiome shifts in high- and low-gastric-cancer-risk populations from Colombia, South America. Gut Microbes 15:2186677.

11. Lapidot Y, Reshef L, Cohen D, Muhsen K. 2021. *Helicobacter pylori* and the intestinal microbiome among healthy school-age children. Helicobacter 26:e12854.

12. Donald K, Finlay BB. 2023. Early-life interactions between the microbiota and immune system: impact on immune system development and atopic disease. Nat Rev Immunol 23:735–748.

13. Keady MM, Jimenez RR, Bragg M, Wagner JCP, Bornbusch SL, Power ML, Muletz-Wolz CR. 2023. Ecoevolutionary processes structure milk microbiomes across the mammalian tree of life. Proc Natl Acad Sci U S A 120:e2218900120.

14. Ganal-Vonarburg SC, Hornef MW, Macpherson AJ. 2020. Microbial-host molecular exchange and its functional consequences in early mammalian life. Science 368:604– 607.

15. Le Doare K, Holder B, Bassett A, Pannaraj PS. 2018. Mother’s milk: A purposeful contribution to the development of the infant Microbiota and immunity. Front Immunol 9:361.

16. Yatsunenko T, Rey FE, Manary MJ, Trehan I, Dominguez-Bello MG, Contreras M, Magris M, Hidalgo G, Baldassano RN, Anokhin AP, Heath AC, Warner B, Reeder J, Kuczynski J, Caporaso JG, Lozupone CA, Lauber C, Clemente JC, Knights D, Knight R, Gordon JI. 2012. Human gut microbiome viewed across age and geography. Nature 486:222–227.

17. Kennedy EA, Weagley JS, Kim AH, Antia A, DeVeaux AL, Baldridge MT. 2025. Microbiota assembly of specific pathogen-free neonatal mice. bioRxiv.

18. Hughes KR, Schofield Z, Dalby MJ, Caim S, Chalklen L, Bernuzzi F, Alcon-Giner C, Le Gall G, Watson AJM, Hall LJ. 2020. The early life microbiota protects neonatal mice from pathological small intestinal epithelial cell shedding. FASEB J 34:7075–7088.

19. Al Nabhani Z, Dulauroy S, Marques R, Cousu C, Al Bounny S, Déjardin F, Sparwasser T, Bérard M, Cerf-Bensussan N, Eberl G. 2019. A weaning reaction to Microbiota is required for resistance to immunopathologies in the adult. Immunity 50:1276–1288.e5.

20. Schwarz S, Morelli G, Kusecek B, Manica A, Balloux F, Owen RJ, Graham DY, van der Merwe S, Achtman M, Suerbaum S. 2008. Horizontal versus familial transmission of *Helicobacter pylori*. PLoS Pathog 4:e1000180.

21. Konno M, Yokota S-I, Suga T, Takahashi M, Sato K, Fujii N. 2008. Predominance of mother-to-child transmission of *Helicobacter pylori* infection detected by random amplified polymorphic DNA fingerprinting analysis in Japanese families. Pediatr Infect Dis J 27:999–1003.

22. Kienesberger S, Perez-Perez GI, Olivares AZ, Bardhan P, Sarker SA, Hasan KZ, Sack RB, Blaser MJ. 2018. When is *Helicobacter pylori* acquired in populations in developing countries? A birth-cohort study in Bangladeshi children. Gut Microbes 9:252–263.

23. Salama NR, Hartung ML, Müller A. 2013. Life in the human stomach: persistence strategies of the bacterial pathogen *Helicobacter pylori*. Nat Rev Microbiol 11:385–399.

24. van Zwet AA, Thijs JC, Kooistra-Smid AM, Schirm J, Snijder JA. 1994. Use of PCR with feces for detection of *Helicobacter pylori* infections in patients. J Clin Microbiol 32:1346–1348.

25. Parsonnet J, Shmuely H, Haggerty T. 1999. Fecal and oral shedding of *Helicobacter pylori* from healthy infected adults. JAMA 282:2240–2245.

26. Li C, Ha T, Ferguson DA Jr, Chi DS, Zhao R, Patel NR, Krishnaswamy G, Thomas E. 1996. A newly developed PCR assay of H. pylori in gastric biopsy, saliva, and feces. Evidence of high prevalence of H. pylori in saliva supports oral transmission. Dig Dis Sci 41:2142–2149.

27. Minoura T, Kato S, Otsu S, Kodama M, Fujioka T, Iinuma K, Nishizono A. 2005. Influence of age and duration of infection on bacterial load and immune responses to *Helicobacter pylori* infection in a murine model. Clin Exp Immunol 139:43–47.

28. Kløve S, Graversen KB, Rasmussen JA, Assis J, Schluter J, Blaser MJ, Andersen SB. 2025. Early-life Helicobacter pylori infection worsens metabolic state in mice receiving a high-fat diet. bioRxiv.

29. RStudio Team. 2023. RStudio: Integrated Development Environment for R. RStudio, PBC., Boston, MA.

30. Austin GI, Park H, Meydan Y, Seeram D, Sezin T, Lou YC, Firek BA, Morowitz MJ, Banfield JF, Christiano AM, Pe’er I, Uhlemann A-C, Shenhav L, Korem T. 2023. Contamination source modeling with SCRuB improves cancer phenotype prediction from microbiome data. Nat Biotechnol 41:1820–1828.

31. Callahan BJ, McMurdie PJ, Rosen MJ, Han AW, Johnson AJA, Holmes SP. 2016. DADA2: High-resolution sample inference from Illumina amplicon data. Nat Methods 13:581–583.

32. Quast C, Pruesse E, Gerken J, Schweer T, Yilmaz P, Peplies J, Glockner FO. 2012. SILVA Databases, p. 1–11. In Encyclopedia of Metagenomics. Springer New York, New York, NY.

33. McMurdie PJ, Holmes S. 2013. phyloseq: an R package for reproducible interactive analysis and graphics of microbiome census data. PLoS One 8:e61217.

34. Oksanen J, Simpson GL, Blanchet FG, Kindt R, Legendre P, Minchin PR, O’Hara RB, Solymos P, Stevens MHH, Szoecs E, Wagner H, Barbour M, Bedward M, Bolker B, Borcard D, Carvalho G, Chirico M, De Caceres M, Durand S, Evangelista HBA, FitzJohn R, Friendly M, Furneaux B, Hannigan G, Hill MO, Lahti L, McGlinn D, Ouellette M-H, Ribeiro Cunha E, Smith T, Stier A, Ter Braak CJF, Weedon J. 2024. vegan: Community Ecology Package.

35. Barnett D, Arts I, Penders J. 2021. microViz: an R package for microbiome data visualization and statistics. J Open Source Softw 6:3201.

36. Cao Y, Dong Q, Wang D, Zhang P, Liu Y, Niu C. 2022. microbiomeMarker: an R/Bioconductor package for microbiome marker identification and visualization. Bioinformatics 38:4027–4029.

37. Chen S, Zhou Y, Chen Y, Gu J. 2018. fastp: an ultra-fast all-in-one FASTQ preprocessor. Bioinformatics 34:i884–i890.

38. Patel H, Ewels P, Manning J, Garcia MU, Peltzer A, Hammarén R, Botvinnik O, Talbot A, Sturm G, Bot N-C, Zepper M, Moreno D, Vemuri P, Binzer-Panchal M, Greenberg E, silviamorins, Pantano L, Syme R, Kelly G, Hanssen F, Fellows Yates JA, Espinosa- Carrasco J, rfenouil, Zappia L, Cheshire C, Miller E, marchoeppner, Zhou P, Guinchard S, Gabernet G. 2024. nf-core/rnaseq: nf-core/rnaseq v3.18.0 - Lithium Lynx. Zenodo. https://zenodo.org/records/14537300. Retrieved 6 February 2025.

39. Dobin A, Davis CA, Schlesinger F, Drenkow J, Zaleski C, Jha S, Batut P, Chaisson M, Gingeras TR. 2013. STAR: ultrafast universal RNA-seq aligner. Bioinformatics 29:15– 21.

40. Patro R, Duggal G, Love MI, Irizarry RA, Kingsford C. 2017. Salmon provides fast and bias-aware quantification of transcript expression. Nat Methods 14:417–419.

41. Love MI, Huber W, Anders S. 2014. Moderated estimation of fold change and dispersion for RNA-seq data with DESeq2. bioRxiv. bioRxiv.

42. Szklarczyk D, Kirsch R, Koutrouli M, Nastou K, Mehryary F, Hachilif R, Gable AL, Fang T, Doncheva NT, Pyysalo S, Bork P, Jensen LJ, von Mering C. 2023. The STRING database in 2023: protein-protein association networks and functional enrichment analyses for any sequenced genome of interest. Nucleic Acids Res 51:D638– D646.

43. Griss J, Viteri G, Sidiropoulos K, Nguyen V, Fabregat A, Hermjakob H. 2020. ReactomeGSA - efficient multi-omics comparative pathway analysis. Mol Cell Proteomics 19:2115–2125.

44. Li X, Zhang C, Gong T, Ni X, Li J, Zhan D, Liu M, Song L, Ding C, Xu J, Zhen B, Wang Y, Qin J. 2018. A time-resolved multi-omic atlas of the developing mouse stomach. Nat Commun 9:4910.

45. Fung C, Tan S, Nakajima M, Skoog EC, Camarillo-Guerrero LF, Klein JA, Lawley TD, Solnick JV, Fukami T, Amieva MR. 2019. High-resolution mapping reveals that microniches in the gastric glands control *Helicobacter pylori* colonization of the stomach. PLoS Biol 17:e3000231.

46. Keilberg D, Zavros Y, Shepherd B, Salama NR, Ottemann KM. 2016. Spatial and temporal shifts in bacterial biogeography and gland occupation during the development of a chronic infection. MBio 7.

47. Takeoka Y, Kataoka K. 1986. Histogenesis of the mouse pyloric mucosa with special reference to the development of surface mucous cells and pylorocytes, and the formation of the generative zone. Arch Histol Jpn 49:519–534.

48. Helander HF. 1970. Gastric acidity in young and adult mice. Scand J Gastroenterol 5:221–224.

49. McConnell EL, Basit AW, Murdan S. 2008. Measurements of rat and mouse gastrointestinal pH, fluid and lymphoid tissue, and implications for in-vivo experiments. J Pharm Pharmacol 60:63–70.

50. Haque M, Koski KG, Scott ME. 2021. A gastrointestinal nematode in pregnant and lactating mice alters maternal and neonatal microbiomes. Int J Parasitol 51:945–957.

51. Depommier C, Everard A, Druart C, Plovier H, Van Hul M, Vieira-Silva S, Falony G, Raes J, Maiter D, Delzenne NM, de Barsy M, Loumaye A, Hermans MP, Thissen J-P, de Vos WM, Cani PD. 2019. Supplementation with *Akkermansia muciniphila* in overweight and obese human volunteers: a proof-of-concept exploratory study. Nat Med 25:1096–1103.

52. Everard A, Belzer C, Geurts L, Ouwerkerk JP, Druart C, Bindels LB, Guiot Y, Derrien M, Muccioli GG, Delzenne NM, de Vos WM, Cani PD. 2013. Cross-talk between *Akkermansia muciniphila* and intestinal epithelium controls diet-induced obesity. Proceedings of the National Academy of Sciences 110:9066–9071.

53. Shin N-R, Lee J-C, Lee H-Y, Kim M-S, Whon TW, Lee M-S, Bae J-W. 2014. An increase in the *Akkermansia spp.* population induced by metformin treatment improves glucose homeostasis in diet-induced obese mice. Gut 63:727–735.

54. Fan S, Jiang Z, Zhang Z, Xing J, Wang D, Tang D. 2023. *Akkermansia muciniphila*: a potential booster to improve the effectiveness of cancer immunotherapy. J Cancer Res Clin Oncol 149:13477–13494.

55. Cox LM, Blaser MJ. 2013. Pathways in microbe-induced obesity. Cell Metab 17:883– 894.

56. Nobel YR, Cox LM, Kirigin FF, Bokulich NA, Yamanishi S, Teitler I, Chung J, Sohn J, Barber CM, Goldfarb DS, Raju K, Abubucker S, Zhou Y, Ruiz VE, Li H, Mitreva M, Alekseyenko AV, Weinstock GM, Sodergren E, Blaser MJ. 2015. Metabolic and metagenomic outcomes from early-life pulsed antibiotic treatment. Nat Commun 6.

57. Trust TJ, Doig P, Emödy L, Kienle Z, Wadström T, O’Toole P. 1991. High-affinity binding of the basement membrane proteins collagen type IV and laminin to the gastric pathogen *Helicobacter pylori*. Infect Immun 59:4398–4404.

58. Rautelin HI, Oksanen AM, Veijola LI, Sipponen PI, Tervahartiala TI, Sorsa TA, Lauhio A. 2009. Enhanced systemic matrix metalloproteinase response in *Helicobacter pylori* gastritis. Ann Med 41:208–215.

59. Dubreuil JD, Giudice GD, Rappuoli R. 2002. *Helicobacter pylori* interactions with host serum and extracellular matrix proteins: potential role in the infectious process. Microbiol Mol Biol Rev 66:617–29, table of contents.

60. Ashraf AA, Gamal SM, Ashour H, Aboulhoda BE, Rashed LA, Harb IA, Abdelfattah GH, El-Seidi EA, Shawky HM. 2021. Investigating *Helicobacter pylori*-related pyloric hypomotility: functional, histological, and molecular alterations. Am J Physiol Gastrointest Liver Physiol 321:G461–G476.

61. Suzuki H, Moayyedi P. 2013. *Helicobacter pylori* infection in functional dyspepsia. Nat Rev Gastroenterol Hepatol 10:168–174.

62. Sýkora J, Malán A, Záhlava J, VarvarsCká J, StoziCcký F, Siala K, Schwarz J. 2004. Gastric emptying of solids in children with *H*. pyloriUpositive and H. pyloriUnegative nonUulcer dyspepsia. J Pediatr Gastroenterol Nutr 39:246–252.

63. Nascakova Z, He J, Papa G, Francas B, Azizi F, Müller A. 2024. *Helicobacter pylori* induces the expression of Lgr5 and stem cell properties in gastric target cells. Life Sci Alliance 7:e202402783.

