## Supplementary Figs+Table for "Neonatal infection with *Helicobacter pylori* affects stomach and colon microbiome composition and gene expression in mice"

**Supplementary figures**

**Fig. S1**

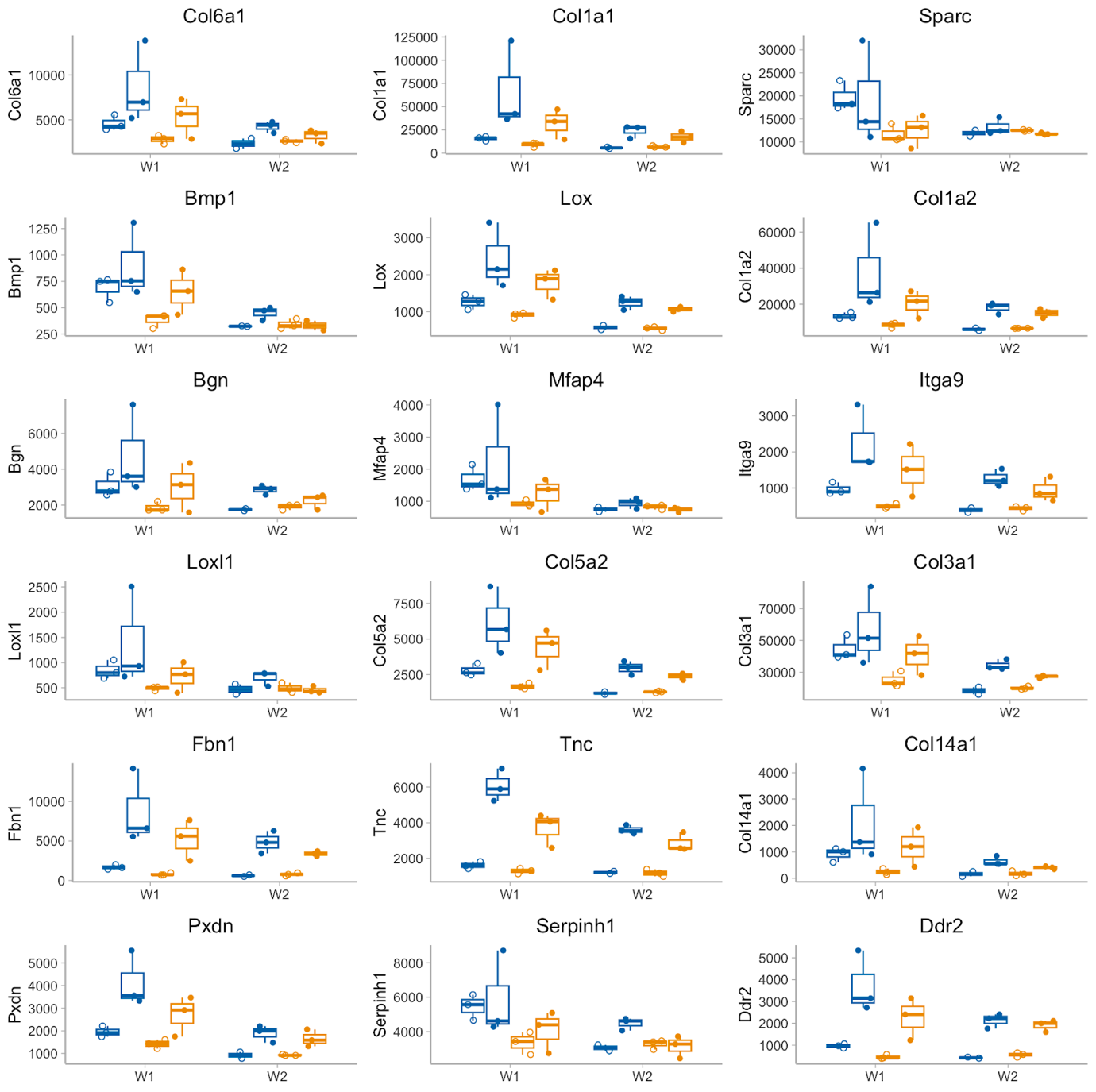

***Fig. S1:*** *Stomach tissue gene expression of genes related to the extracellular matrix from samples collected one week after gavage (W1) and two weeks after gavage (W2) from mice infected with H. pylori (yellow) or treated with a control solution (blue). Expression levels varied between sequencing runs (open circles = Novogene, filled circles = Rigshospitalet). However, for the majority of genes the pattern between treatment was the same, with overall higher expression levels in the Rigshospitalet data set. Note the different scales on the y-axis between plots.*

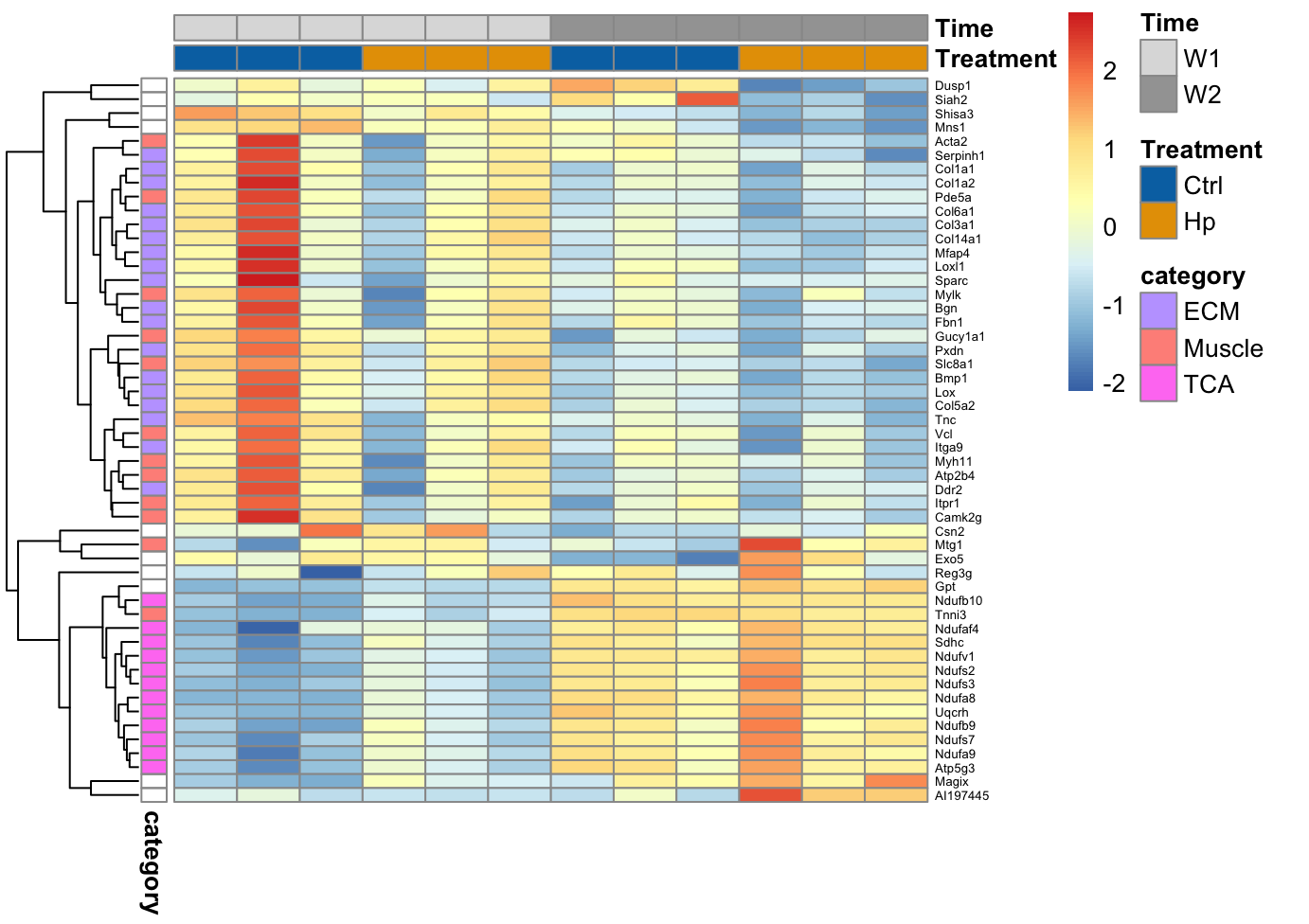

***Fig. S2:*** *Heatmap of scaled expression of genes that were differentially expressed with a p_adj_ < 0.05, or 0.1 and classified as belonging to an enriched Reactome pathway in the STRING analyses (ECM = extracellular matrix, Muscle = muscle contraction, and TCA = the citric acid cycle). Only data from Rigshospitalet is shown, due to the differences in expression levels between sequencing runs (see* ***Fig. 10*** *for the Novogene dataset).*

**Table S1**

*Overview of mice included in the study and indication of sampling time relative to last infection with H. pylori (Hp) or control (Ctrl), and relative to weaning as well as indications of what analyses have been applied****.*** *For mice sacrificed before weaning, sex was estimated after the RNAseq analysis based on Xist gene expression, which is only expressed in females.*

| **Sampling time** | |  |  |  |  |  |
| --- | --- | --- | --- | --- | --- | --- |
| **Time since**  **last infection** | **Time since**  **weaning** | **Treatment** | **Litter** | **Sex** | **Stomach and SI Hp quantification by qPCR** | **Stomach and colon 16S seq and stomach RNAseq** |
| 2 days | NA | **Hp** | 4a | NA | Yes | - |
| 2 days | NA | **Hp** | 4a | NA | Yes | - |
| 2 days | NA | **Hp** | 4a | NA | Yes | - |
| 2 days | NA | **Hp** | 4a | NA | Yes | - |
| 2 days | NA | **Hp** | 4a | NA | Yes | - |
| 2 days | NA | **Hp** | 4a | NA | Yes | - |
| 1 week | NA | **Hp** | 6a | F | Yes | Yes |
| 1 week | NA | **Hp** | 6a | M | Yes | Yes |
| 1 week | NA | **Hp** | 6b | F | Yes | Yes |
| 1 week | NA | **Hp** | 15a | M | Yes | Yes |
| 1 week | NA | **Hp** | 15a | M | Yes | Yes |
| 1 week | NA | **Hp** | 15b | M | Yes | Yes |
| 1 week | NA | Ctrl | 14a | F | Yes / neg control | Yes |
| 1 week | NA | Ctrl | 14a | F | - | Yes |
| 1 week | NA | Ctrl | 14a | F | - | Yes |
| 1 week | NA | Ctrl | 14b | F | - | Yes |
| 1 week | NA | Ctrl | 14b | F | - | Yes |
| 1 week | NA | Ctrl | 14b | M | - | Yes |
| 2 weeks | NA | **Hp** | 6a | M | Yes | Yes |
| 2 weeks | NA | **Hp** | 6a | M | Yes | Yes |
| 2 weeks | NA | **Hp** | 6b | F | Yes | Yes |
| 2 weeks | NA | **Hp** | 15a | M | Yes | Yes |
| 2 weeks | NA | **Hp** | 15a | F | Yes | Yes |
| 2 weeks | NA | **Hp** | 15b | M | Yes | Yes |
| 2 weeks | NA | Ctrl | 14a | M | Yes / neg control | Yes |
| 2 weeks | NA | Ctrl | 14a | M | - | Yes |
| 2 weeks | NA | Ctrl | 14a | F | - | Yes |
| 2 weeks | NA | Ctrl | 14b | NA | - | Yes, but RNAseq failed |
| 2 weeks | NA | Ctrl | 14b | M | - | Yes |
| 2 weeks | NA | Ctrl | 14b | F | - | Yes |
| 3 weeks | 2 days | **Hp** | 15a | F | Yes | - |
| 3 weeks | 2 days | **Hp** | 6b | F | Yes | - |
| 3 weeks | 2 days | **Hp** | 6b | F | Yes | - |
| 3 weeks | 2 days | **Hp** | 15a | M | Yes | - |
| 3 weeks | 2 days | **Hp** | 6a | M | Yes | - |
| 3 weeks | 2 days | **Hp** | 6a | M | Yes | - |
| 4 weeks | 10 days | **Hp** | 15b | F | Yes | - |
| 4 weeks | 10 days | **Hp** | 6a | F | Yes | - |
| 4 weeks | 10 days | **Hp** | 15a | M | Yes | - |
| 4 weeks | 10 days | **Hp** | 15b | M | Yes | - |
| 4 weeks | 10 days | **Hp** | 6b | M | Yes | - |
| 4 weeks | 10 days | **Hp** | 6b | M | Yes | - |
